# Molecular detection of pathogens in ticks and fleas collected from companion dogs and cats in East and Southeast Asia

**DOI:** 10.1101/2020.05.27.118554

**Authors:** Viet-Linh Nguyen, Vito Colella, Grazia Greco, Fang Fang, Wisnu Nurcahyo, Upik Kesumawati Hadi, Virginia Venturina, Kenneth Boon Yew Tong, Yi-Lun Tsai, Piyanan Taweethavonsawat, Saruda Tiwananthagorn, Sahatchai Tangtrongsup, Thong Quang Le, Khanh Linh Bui, Thom Do, Malaika Watanabe, Puteri Azaziah Megat Abd Rani, Filipe Dantas-Torres, Lenaig Halos, Frederic Beugnet, Domenico Otranto

## Abstract

Ticks and fleas are considered amongst the most important arthropod vectors of medical and veterinary concern due to their ability to transmit pathogens to a range of animal species including dogs, cats and humans. By sharing a common environment with humans, companion animal-associated parasitic arthropods may potentially transmit zoonotic vector-borne pathogens (VBPs). This study aimed to molecularly detect pathogens from ticks (*n* = 392) and fleas (*n* = 248) collected from companion dogs and cats in East and Southeast Asia. Of the 392 ticks tested, 37 (9.4%) scored positive for at least one pathogen with *Hepatozoon canis* being the most prevalent (5.4%), followed by *Ehrlichia canis* (1.8%), *Babesia vogeli* (1%), *Anaplasma platys* (0.8%) and *Rickettsia* spp. (1%) [including *Rickettsia* sp. (0.5%), *Rickettsia asembonensis* (0.3%), *Rickettsia felis* (0.3%)]. Out of 248 fleas tested, 106 (42.7%) were harboring at least one pathogen with *R. felis* being the most common (19.4%), followed by *Bartonella* spp. (16.5%), *Rickettsia asembonensis* (10.9%) and *Candidatus* Rickettsia senegalensis (0.4%). Furthermore, 35 *Rhipicephalus sanguineus* sensu lato ticks were subjected to phylogenetic analysis, of which 34 ticks belonged to the tropical and only one belonged to the temperate lineage (*Rh. sanguineus* sensu stricto). Our data reveals the circulation of different VBPs in ticks and fleas of dogs and cats from Asia, including zoonotic agents, which may represent a potential risk to animal and human health.

**Author summary:** Ticks and fleas are among the most important vectors of pathogens infesting many animal species including humans worldwide. Although a number of vector-borne pathogens have been detected in dogs and cats from East and Southeast Asia, investigation in ticks and fleas collected from them are scant. In order to provide an overview of the pathogens circulating in ticks and fleas from companion dogs and cats in Asia, ticks (*n* = 392) and fleas (*n* = 248) were collected from privately-owned dogs and cats from China, Taiwan, Indonesia, Malaysia, Singapore, Thailand, Philippines and Vietnam and molecularly screened for the presence of pathogens. Overall, multiple pathogens were found in ticks (i.e., *Babesia vogeli*, *Hepatozoon canis*, *Ehrlichia canis*, *Anaplasma platys* and *Rickettsia* spp.) and fleas (i.e., *Rickettsia* spp. and *Bartonella* spp.) from the sampling areas. Of the ticks tested, 9.4% scored positive for at least one pathogen and of fleas 42.7% harbored at least one pathogen with *Rickettsia felis* being the most common (19.4%). Overall, of the detected pathogens, *R. felis* stood out as the most important due to its zoonotic potential. The result of this study should increase awareness among pet owners and veterinary practitioners regarding the importance of ticks and fleas, and their transmitted pathogens.

## Introduction

Vector-borne diseases are caused by bacteria, viruses, protozoa and helminths transmitted by arthropod vectors, including ticks and fleas, worldwide [1]. For instance, *Rhipicephalus sanguineus* sensu lato ticks play an important role in the transmission of many pathogens to dogs (e.g., *Ehrlichia canis*, *Rickettsia conorii*, *Rickettsia rickettsia*, *Babesia vogeli* and *Hepatozoon canis*), some of which may also infect humans [2–4]. The cat flea *Ctenocephalides felis* is the main vector of *Bartonella henselae*, the main causative agent of cat-scratch disease to humans [5–6] and is also considered as the main vector of *Rickettsia felis* [7].

East (EA) and Southeast Asia (SEA) are among the world’s fastest-growing economic regions [8], which also resulted in a rise of populations of companion dogs and cats [9]. Companion dogs and cats live in close association with humans, potentially carrying ticks and fleas into human settlements. A large-scale survey conducted in EA and SEA reported that 22.3% of dogs and 3.7% of cats were infested by ticks, while 14.8% of dogs and 19.6% of cats were infested by fleas [10]. The most common flea species parasitizing dogs and cats in EA and SEA is *C. felis*, with *Ctenocephalides orientis* being increasingly observed in dogs [10–11]. *Rhipicephalus sanguineus* s.l., *Rhipicephalus haemaphysaloides* and *Haemaphysalis longicornis* represent the most common tick species reported in dogs and cats [10–14]. These tick species are responsible for the transmission of several species of apicomplexan protozoan of the genus *Babesia. Babesia vogeli* was reported in cats in Thailand and China [15–16] and widely reported in dogs in EA and SEA, including China, Indonesia, Cambodia, Thailand, Malaysia and Philippines [17–20]. Additionally, dogs from Taiwan, Malaysia, China, and Singapore [10,21,22] were also diagnosed with *Babesia gibsoni* infection. Other apicomplexan parasites commonly found in dogs across this region are those of the species *H. canis* transmitted by ingestion of *Rh. sanguineus* s.l. This protozoan is commonly found in dogs in Thailand, Taiwan, China, Cambodia, Malaysia, Philippines and Vietnam [10,17–19,23–25] and in cats in Thailand and Philippines [10,23]. Of the tick-borne anaplasmatacea bacteria, *Anaplasma platys* was found in dogs from Cambodia [18] and cats from Thailand [26]. Apart from tick-borne pathogens, flea-borne pathogens are also increasingly recognized as important pathogenic agents to animals and humans. For instance, *R. felis*, the etiological agent of flea-borne spotted fever in humans, has been detected in dogs and in *C. felis* from Taiwan, Laos, Malaysia, China and Cambodia with the prevalence ranging from 10 to 60% [23,27–29]. Other zoonotic flea-borne pathogens such as *Bartonella clarridgeiae* and *B. henselae*, agents of which cause cat scratch disease, were molecularly detected in cats and their fleas from Indonesia, Singapore, Philippines, Thailand, Malaysia and China [27,30–34].

Despite many scientific investigations reported the circulation of vector-borne pathogens (VBPs) in dogs and cats in EA and SEA, there is a lack of studies conducted with their associated ticks and fleas, especially those of zoonotic importance. Therefore, the current study aimed to provide an overview of the pathogens circulating in ticks and fleas from companion dogs and cats in EA and SEA.

## Materials and methods

### Ethics approval and consent to participate

The protocol of this study was approved by the Ethical Committee of the Department of Veterinary Medicine of the University of Bari (Prot. no. 13/17). All animals’ owners have read, approved and signed an owner informed consent which contained information on study procedures and aims of this study.

### Samples collection and DNA isolation

Of the 2381 privately-owned animals examined (i.e., 1229 dogs and 1152 cats), ticks and fleas were collected from 401 infested animals (i.e., 271 dogs and 130 cats) from China, Taiwan, Indonesia, Malaysia, Singapore, Thailand, Philippines and Vietnam under the context of a previous multicenter survey [10]. Ticks and fleas were collected and placed in labeled tubes individualized per host, containing 70% ethanol. Ticks and fleas (20%) were randomly selected from each tick/flea species and from each infested animal in all studied countries, giving a total number of 392 ticks (i.e., 377 *Rh. sanguineus* s.l., three *Rh. haemaphysaloides*, seven *H. longicornis*, two *Haemaphysalis wellingtoni*, one *Haemaphysalis hystricis*, one *Haemaphysalis campanulata* and one *Ixodes* sp.) from 248 animals (39 cats and 209 dogs) (Table 1) and 248 fleas (i.e., 209 *C. felis*, 38 *C. orientis* and one *Xenopsylla cheopis*) from 213 animals (104 cats and 109 dogs) were subjected to DNA isolation individually (Table 2). Data on the molecular identification of these ticks and fleas are available elsewhere (see Table 4 of Ref. [10]) Genomic DNA was isolated according to the procedures previously described [35].

**Table 1.**
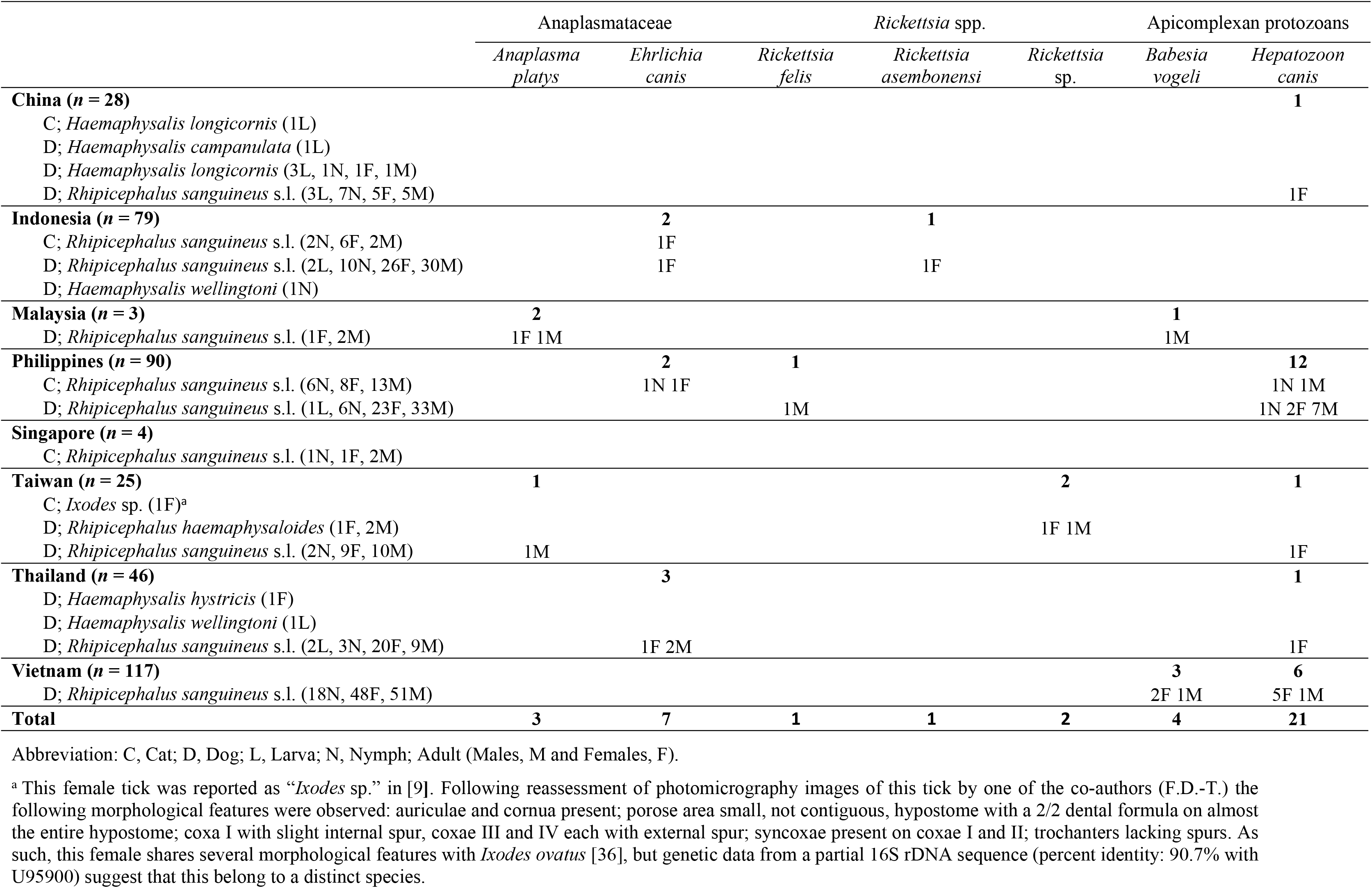
Pathogens detected in ticks according to their species, developmental stage, sex and host in East and Southeast Asia.

**Table 2.** Pathogens detected in fleas according to their species, developmental stage, sex and host in East and Southeast Asia.

|  | *Rickettsia* spp. |  |  | *Bartonella* spp. |
| --- | --- | --- | --- | --- |
|  | *Rickettsia felis* | *Rickettsia asembonensis* | *Candidatus Rickettsia senegalensis* |  |
| **China (n = 17)** | **1** |  |  | **2** |
| C; *Ctenocephalides felis* (8F, 2M) | 1F |  |  | 2F |
| D; *Ctenocephalides felis* (7F) |  |  |  |  |
| **Indonesia (n = 48)** | **12** | **9** |  | **21** |
| C; *Ctenocephalides felis* (4F, 24M) | 7F 2M |  |  | 18F |
| C; *Xenopsylla cheopis* (1M) |  |  |  | 1M |
| D; *Ctenocephalides felis* (6F, 1M) | 2F |  |  | 2F |
| D; *Ctenocephalides orientis* (9F, 3M) | 1M | 7F 2M |  |  |
| **Malaysia (n = 7)** |  |  |  |  |
| C; *Ctenocephalides felis* (4F, 1M) |  |  |  |  |
| D; *Ctenocephalides felis* (2F) |  |  |  |  |
| **Philippines (n = 90)** | **20** | **9** |  | **4** |
| C; *Ctenocephalides felis* (24F, 9M) | 4F 3M |  |  | 3F 1M |
| D; *Ctenocephalides felis* (34F, 13M) | 12F 1M | 1F |  |  |
| D; *Ctenocephalides orientis* (9F, 1M) |  | 8F |  |  |
| **Singapore (n = 2)** |  |  |  | **2** |
| C; *Ctenocephalides felis* (1M) |  |  |  | 1M |
| C; *Ctenocephalides orientis* (1F) |  |  |  | 1F |
| **Taiwan (n = 24)** | **8** |  |  | **4** |
| C; *Ctenocephalides felis* (10F, 4M) | 3F 2M |  |  | 3F 1M |
| D; *Ctenocephalides felis* (6F, 3M) | 3F |  |  |  |
| D; *Ctenocephalides orientis* (1F) |  |  |  |  |
| **Thailand (n = 21)** | **1** | **4** | **1** | **5** |
| C; *Ctenocephalides felis* (4F, 1M) |  |  |  | 2F |
| D; *Ctenocephalides felis* (7F, 4M) | 1F |  |  | 1F |
| D; *Ctenocephalides orientis* (2F, 3M) |  | 2F 2M | 1M | 1F 1M |
| **Vietnam (n = 39)** | **6** | **5** |  | **3** |
| C; *Ctenocephalides felis* (14F, 5M) | 3F 1M |  |  | 3F |
| D; *Ctenocephalides felis* (8F, 3M) | 2F |  |  |  |
| D; *Ctenocephalides orientis* (7F, 2M) |  | 4F 1M |  |  |
| **Total** | **48** | **27** | **1** | **41** |

Abbreviation: C, Cat; D, Dog; F, Female; M, Male.

### Molecular detection and phylogenetic analysis of pathogens

Tick DNA samples were tested for the presence of apicomplexan protozoans (i.e., *Babesia* spp., *Hepatozoon* spp.), Anaplasmataceae (i.e., *Anaplasma* spp., *Ehrlichia* spp.) and *Coxiella burnetii* by conventional PCR (cPCR). Flea DNA samples were tested by using real-time PCR for *Bartonella* spp. The occurrence of *Rickettsia* spp. was detected in both tick and flea samples. In particular, the first cPCR amplified a portion of citrate synthase (*gltA*) gene, which is presented in all members of the genus *Rickettsia*. Positive samples were then subjected to a second cPCR, which amplified a fragment of the outer membrane protein (*ompA*) of the spotted fever group rickettsiae (SFGR). All primers and PCR protocols used for the detection of VBPs are summarized in Table 3. For all reactions, DNA of pathogen-positive samples served as a positive control. Amplified cPCR products were examined on 2% agarose gels stained with GelRed (VWR International PBI, Milan, Italy) and visualized on a GelLogic 100 gel documentation system (Kodak, New York, USA). The cPCR amplicons were sequenced using the Big Dye Terminator v.3.1 chemistry in a 3130 Genetic analyzer (Applied Biosystems, California, USA). Nucleotide sequences were edited, aligned and analyzed using the BioEdit 7.0 software and compared with those available in the GenBank database using Basic Local Alignment Search Tool (http://blast.ncbi.nlm.nih.gov/Blast.cgi).

**Table 3.**
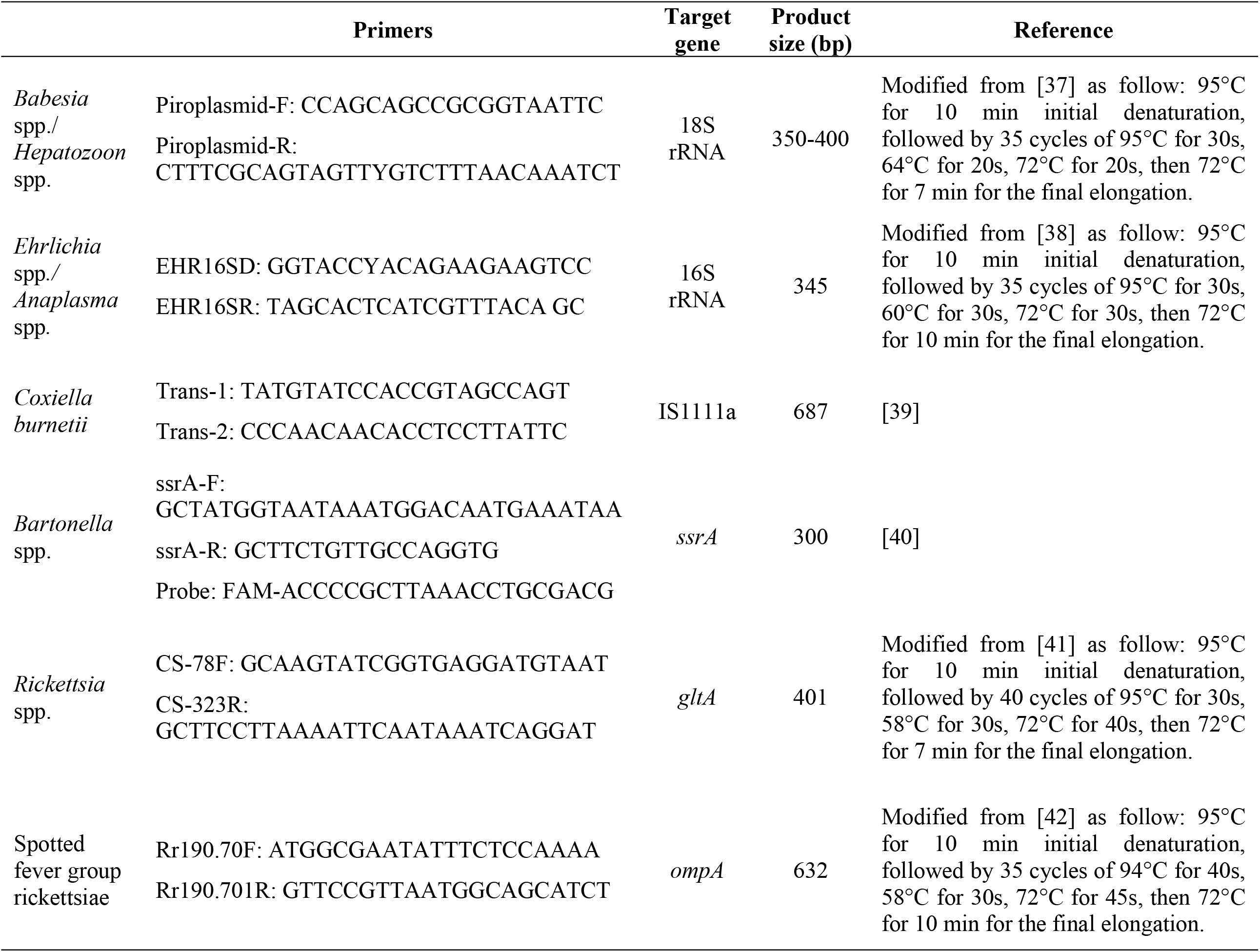
Primers, target genes and PCR conditions used in this study.

To assess the genetic variation of *Rh. sanguineus* s.l. and *Rickettsia* spp., the mitochondrial 16S rDNA sequences of *Rh. sanguineus* s.l. ticks generated previously [10] as well as the *gltA* and *ompA* gene sequences of *Rickettsia* spp. generated herein were subjected to phylogenetic analysis. Phylogenetic relationship was inferred by Maximum Likelihood (ML) method after selecting the best-fitting substitution model. Evolutionary analysis was conducted on 8000 bootstrap replications using the MEGA 7 software [43].

### Statistical analysis

The percentage of detected pathogens was calculated and 95% confidence intervals (by modified Wald method) were estimated by using Quantitative Parasitology 3.0 software [44]. The Fisher’s exact test was performed to analyze statistically significant differences in the detection of pathogens in fleas and ticks, and in the distribution of different *Rickettsia* spp. in flea species using SPSS 16.0 software. Differences were considered significant at *p* < 0.05.

## Results

The occurrence of VBPs has been detected in ticks and fleas, with a higher number of fleas in which at least one pathogen was detected compared to ticks (Fisher’s exact test, *p* < 0.001). Of the 392 ticks tested, 37 (9.4%; 95% CI: 6.9–12.8%) scored positive for at least one pathogen with *H. canis* being the most prevalent (5.4%; 95% CI: 3.5–8.1%), followed by *E. canis* (1.8%; 95% CI: 0.8–3.7%), *B. vogeli* (1%; 95% CI: 0.3–2.7%), *Rickettsia* spp. (1%; 95% CI: 0.3–2.7%) and *A. platys* (0.8%; 95% CI: 0.2–2.3%). Co-infection of *A. platys* and *B. vogeli* was found in one *Rh. sanguineus* s.l., whereas none of the ticks tested positive for *C. burnetii* (Table 1). Out of 248 fleas tested, 106 (42.7%; 95% CI: 36.7–49.0%) were harboring at least one pathogen with *R. felis* being the most common (19.4%; 95% CI: 14.9–24.8%) followed by *Bartonella* spp. (16.5%; 95% CI: 12.4–21.7%), *Rickettsia asembonensis* (10.9%; 95% CI: 7.6–15.4%) and *Candidatus* Rickettsia senegalensis (0.4%; 95% CI: <0.0001–2.5%) (Table 2).

Representative nucleotide sequences for each detected pathogen displayed 99.4-100% identity with those available in GenBank database. In particular, *A. platys* nucleotide sequences (*n* = 3) revealed 99.6-100% identity with KU500914, *H. canis* (*n* = 20) 99.7-100% identity with DQ519358, *E. canis* (*n* = 7) and *B. vogeli* (*n* = 4) 100% identical to MN227484 and KX082917, respectively.

For *Rickettsia* spp. detection in ticks, the *gltA* sequence identified in one tick showed 99.7% identity with *R. asembonensis* (GenBank: KY445723) and another was identical to *R. felis* (100% nucleotide identity with MG845522 and 99.7% nucleotide identity with *R. felis* strain URRWXCal2; GenBank: CP000053). The *ompA* gene amplification was successful for two samples which showed 100% nucleotide identity with unidentified *Rickettsia* sp. (GenBank: EF219467), which is related (98.3%) to *Rickettsia rhipicephali* (GenBank: U43803).

For *Rickettsia* spp. detection in fleas, amplification of a portion of the *gltA* gene was positive from 76 fleas. The partial *ompA* gene was successfully amplified from 17 of the 76 *gltA*-positive *C. felis* fleas. All of those *ompA* gene sequences were 100% identical to *R. felis* strain URRWXCal2 (GenBank: CP000053). Sequence analysis of the *gltA* genes fragment from the other 59 *Rickettsia* positive *C. felis* fleas revealed that 31 sequences had 99.4% nucleotide similarity with *R. felis* strain URRWXCal2 (GenBank: CP000053), 27 sequences 99.7% with *R. asembonensis* (GenBank: KY445723) and one 100% with *Candidatus* R. senegalensis (GenBank: MK548197).

Representative sequences of pathogens detected in this study were deposited in the GenBank database under accession number XXX (AN will be provided in R1).

The phylogenetic tree based on the partial *ompA* gene sequences showed that all *R. felis* isolated from fleas were assembled together in one cluster, whereas *Rickettsia* sp. isolated from ticks clustered with *Rh. rhipicephali* and *Rickettsia massiliae* (Fig 1). In the *gltA* tree, phylogenetical analysis revealed that *R. felis*, *R. asembonensis* and *Candidatus* R. senegalensis herein detected were formed together in a well-supported sister cluster include other *R. felis*-like organisms (RFLO), close to the cluster of *Rickettsia australis* and *Rickettsia akari* (Fig 2).

**Fig 1.**
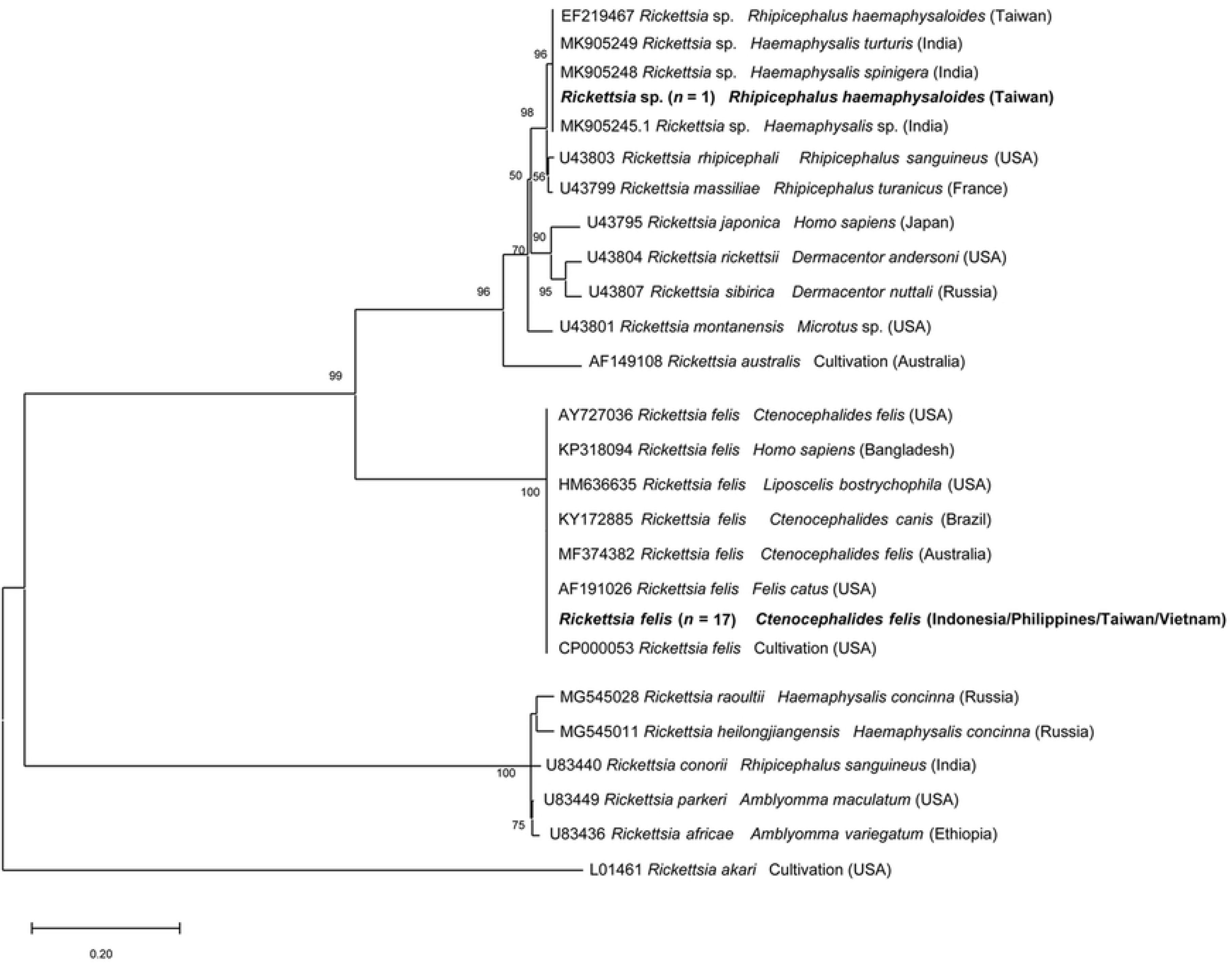
Phylogenetic relationship of *Rickettsia* spp. sequences isolated in this study to other *Rickettsia* spp. based on a partial sequence of the *ompA* gene. The analyses were performed using a Maximum Likelihood method with Tamura 3-parameter model. A discrete Gamma distribution was used to model evolutionary rate differences among sites. Sequences are presented by GenBank accession number, host species and country of origin.

**Fig 2.**
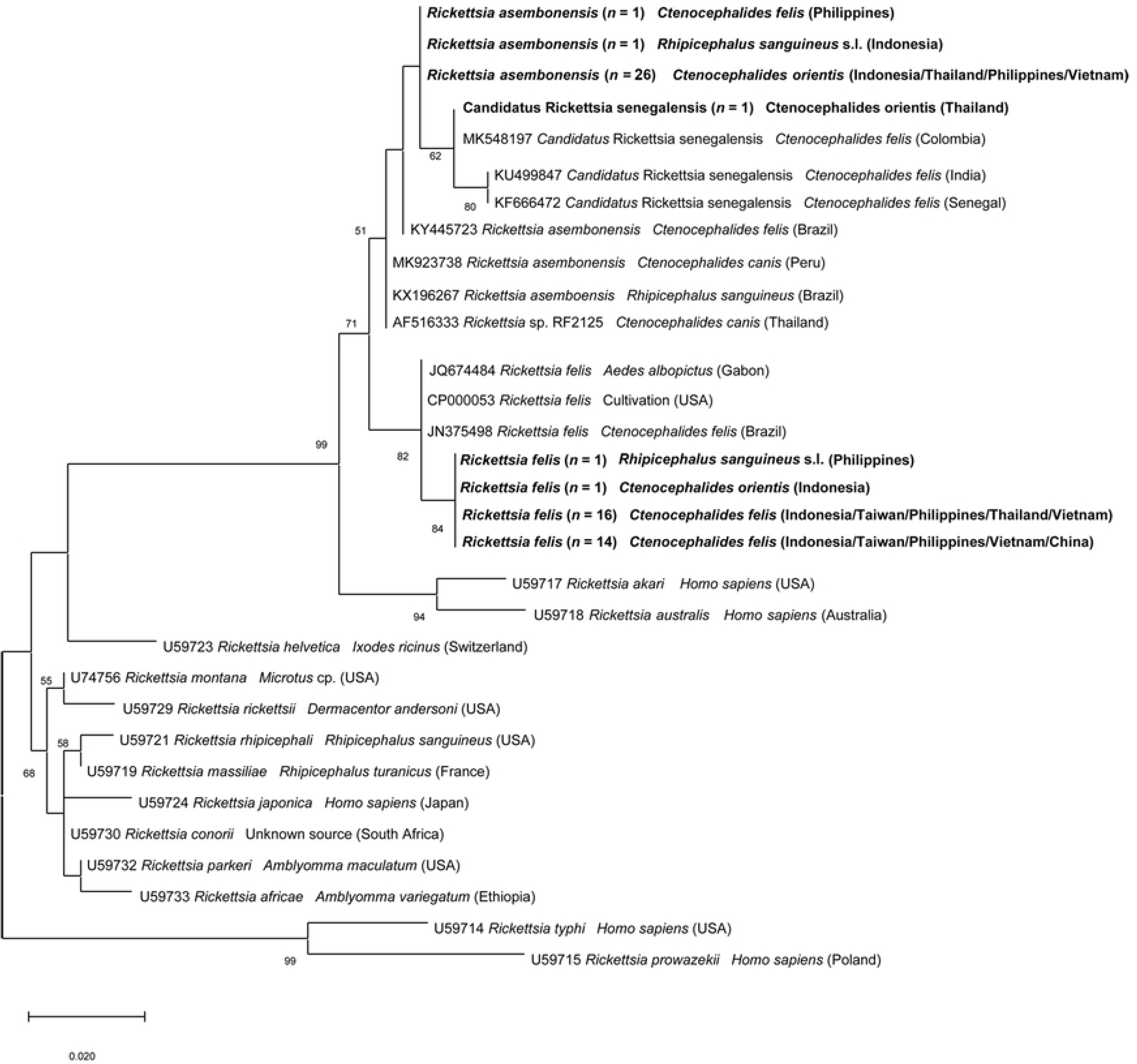
Phylogenetic relationship of *Rickettsia* spp. sequences isolated in this study to other *Rickettsia* spp. based on a partial sequence of the *gltA* gene. The analyses were performed using a Maximum Likelihood method with Tamura 3-parameter model. A discrete Gamma distribution was used to model evolutionary rate differences among sites. Sequences are presented by GenBank accession number, host species and country of origin.

The ML tree of 35 representative sequences of *Rh. sanguineus* s.l. mitochondrial 16S rDNA gene showed that 34 sequences were identical to each other and identified as belong to the tropical lineage of *Rh. sanguineus* s.l. (100% identity with GU553075). One sequence from a tick collected from dog in Beijing (northeast China) clustered in the temperate lineage (100% identity with GU553078), that is, *Rh. sanguineus* sensu stricto (Fig 3).

**Fig 3.**
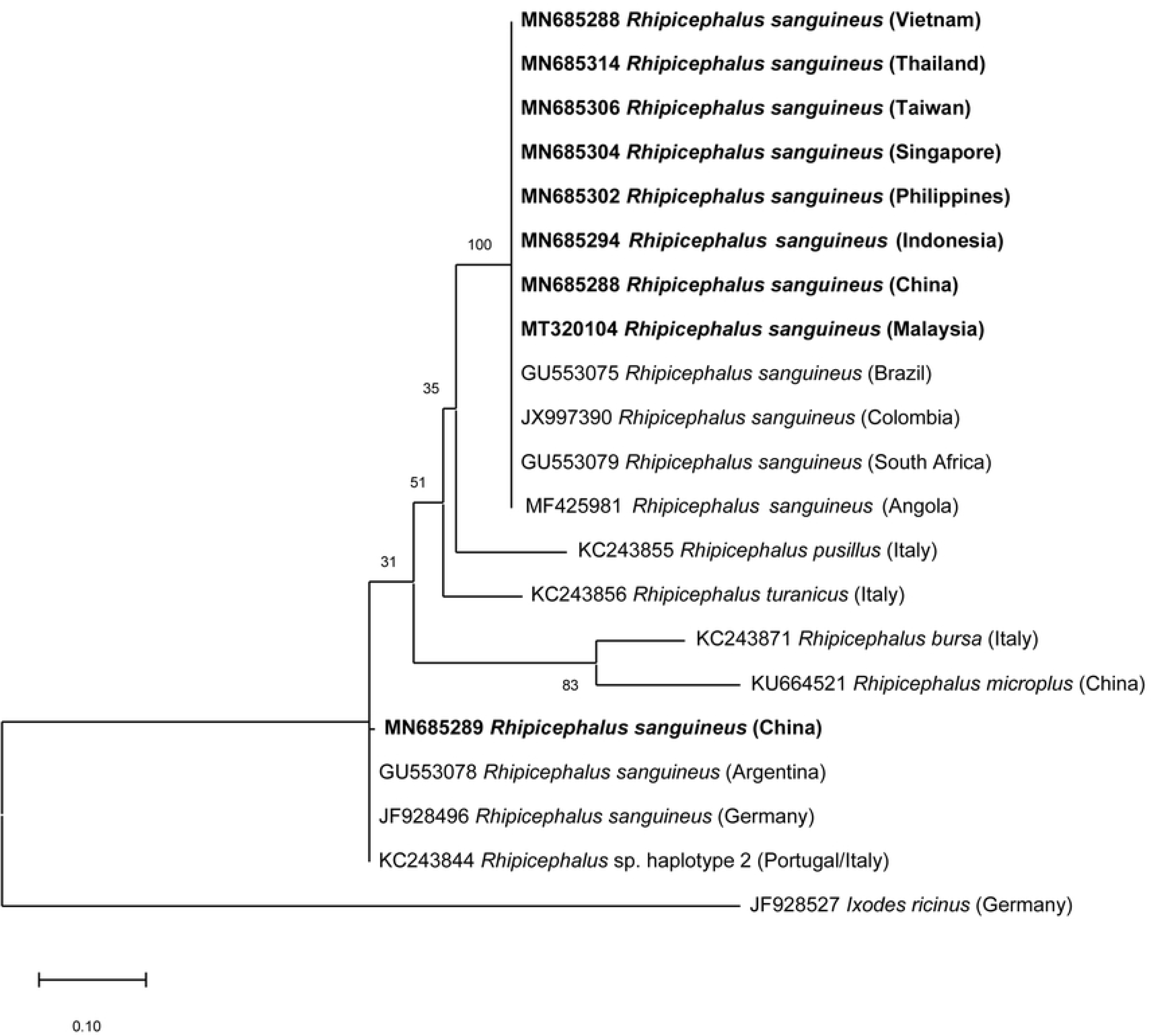
Phylogenetic relationship of the represent *Rhipicephalus sanguineus* s.l. sequences to other *Rhipicephalus* spp. based on a portion of the mitochondrial 16S rDNA sequences. As most sequences (*n* = 34) were identified as belonging to the tropical lineage *Rh. sanguineus* s.l., representatives were selected for each country. The analyses were performed using a Maximum Likelihood method with Tamura 3-parameter model. A discrete Gamma distribution was used to model evolutionary rate differences among sites. Sequences are presented by GenBank accession number, host species and country of origin.

## Discussion

The results of this study reveal the presence of several pathogens in ticks (e.g., *A. platys*, *B. vogeli, E. canis*, *H. canis* and *Rickettsia* spp.) and fleas (e.g., *Rickettsia* spp. and *Bartonella* spp.) collected from dogs and cats in EA and SEA. The relatively low occurrence of pathogens herein detected in ticks is consistent with previous surveys conducted in ticks infesting owned dogs in Asia [13,25,45–47]. Conversely, the occurrence of VBPs is higher in ticks collected from stray animals [23]. Although *B. gibsoni* was found in dogs from China (2.3%; [10]), none of the tested ticks from these dogs was found positive for this parasite. The absence of *B. gibsoni* in tick populations is probably due to the low number of *H. longicornis* and *H. hystricis*, which are recognized as vector of this pathogen [48–49]. The infection of *E. canis* (14.8% by serology) and *H. canis* (1.6% by cPCR) in host populations [10], along with the detection of these pathogens in *Rh. sanguineus* s.l. in the sampling areas support the vector role of this tick species in the transmission of these VBPs in this region [11–14]. The finding of *A. platys* in dogs (7.1% by serology; [10]) and in *Rh. sanguineus* s.l. further suggests its long-suspected, but yet unproven, vector competence for this pathogen. Additionally, the detection of *R. felis* and *R. asembonensis* in *Rh. sanguineus* s.l. is similar to previous results in Chile [50], Brazil [51] and Malaysia [52], consequently giving more concern about the role of *Rh. sanguineus* s.l. in the transmission of *Rickettsia* spp., other than *Rh. conorii*, *Rh. massiliae* and *Rh. rickettsii* [4,53]. *Rickettsia* sp. sequences herein obtained from *Rh. haemaphysaloides* are identical to those previously generated from the same tick species in Taiwan (named *Rickettsia* sp. TwKM01) [54]. This genotype and its closest related species *Rh. Rhipicephali* remain of unknown pathogenicity to mammals [54,55]. Additionally, the vector role of *Rh. haemaphysaloides* needs further investigations since it was found harboring multiple pathogens such as *Rh. rhipicephali*, *A. platys*, *E. canis*, *B. gibsoni* [13,56].

Of the detected VBPs, *R. felis* stood out as the most important due to its worldwide distribution, association with various arthropods, and is recognized as an emerging zoonotic pathogen [57]. In Asia, *R. felis* was first reported in human in Thai-Myanmar border region [58], since then several cases of human infection have been documented in Taiwan [59], Thailand [60], Laos [61], Vietnam [62] and Indonesia [63]. Although *R. felis* was found in many arthropods, including non-hematophagous insect (i.e., the booklouse *Liposcelis bostrychophila*) [64], the widespread of this rickettsia is highly affiliated to the distribution of *C. felis* [57]. *Ctenocephalides felis* is the most well-recognized vector of this rickettsia which transmits the pathogen transovarially and transstadially [65] with dogs as proven mammalian reservoir hosts [66]. The high occurrence of *R. felis* in *C. felis* along with the high relative frequency of occurrence of this flea species in host populations (65.1% in dogs and 98.7% in cats) [10] emphasizes the risk of *R. felis* infection in animals and humans.

In this study, *R. felis* was mostly detected in *C. felis*, whereas *C. orientis* mainly harbored *R. asembonensis* (Fisher’s exact test, *p* < 0.001). The detection of *R. asembonensis* only from fleas collected on dogs (mainly *C. orientis* but in one case in *C. felis*) may suggest that dogs could act as amplifying hosts of this rickettsia, as they do for *R. felis* [66]. *Rickettsia asembonensis* is the most well-characterized genotype of RFLOs [67]. This rickettsia was initially described in fleas from dogs and cats in Kenya [68] and was then reported in various arthropods worldwide [67]. In Asia, *R. asembonensis* was also found in *C. orientis* from dogs [52] and in macaques from Malaysia [69]. Additionally, *Rickettsia* sp. RF2125, a genotype highly related to *R. asembonensis*, was reported with high incidence in *C. orientis* from India and Thailand [70,71], and was also found in a febrile patient from Malaysia [72]. Moreover, *R. felis*, *R. asembonensis* and *Candidatus* R. senegalensis clustered in the SFGR (Fig 3), and while *R. felis* is a recognized pathogen [65], the pathogenicity of other RFLOs is currently unknown.

Besides acting as vectors of *Rickettsia* spp., fleas have been well-recognized as vectors of *Bartonella* spp. [6,73]. The occurrence of *Bartonella*-positive fleas in our study was slightly lower than previous investigations in Thailand [29] Malaysia [33] and Laos [74]. Nevertheless, the occurrence of the two common *Bartonella* spp. (i.e., *B. clarridgeiae* and *B. henselae*) in dogs and cats from EA and SEA is relatively high; up to 60% [18,27,29,75]. Additionally, *B. henselae* infection in humans is usually associated to previous exposure to cats or cat fleas [76], emphasizing the role of cat fleas in *Bartonella* spp. transmission between animals and humans.

In the current study, all tested tick specimens were negative for *C. burnetii* although this pathogen was detected in *Rh. sanguineus* s.l. from dogs in Malaysia [77], in dogs in Taiwan [78] and humans from Thailand [79]. Additionally, *Coxiella*-like symbiont is strongly associated with *Rh. sanguineus* s.l. tropical lineage [80], the role of this microbial in the biology and vectoral capacity of this tick lineage needs further investigation.

Finally, the genetic lineage of *Rh. sanguineus* from EA and SEA was investigated based on the 16S rDNA sequences. The finding of *Rh. sanguineus* s.s., the temperate lineage of *Rh. sanguineus* in Beijing, a cold area, along with the existence of the tropical lineage from all other countries, in warmer localities, agreed with previous studies, which indicated that the tropical lineage is present in areas with annual mean temperature > 20°C, whereas the temperate lineage occurs in areas with annual mean temperature < 20°C [81]. This information is also relevant from a pathogen transmission perspective, considering that different *Rh. sanguineus* s.l. lineages are likely to transmit different pathogens; for instance, *E. canis* is primarily vectored by *Rh. sanguineus* s.l. tropical lineage [82].

Data herein reported updates the list of pathogens occurring in ticks and fleas from companion dogs and cats in EA and SEA. By sharing the common environment with humans, these parasitic arthropods could be responsible for the transmission of pathogens to humans (i.e., *R. felis*). Strategies to prevent tick and flea infestations in these animals are fundamental to decrease the risk of transmission of VBPs to animals and humans.

## Acknowledgments

The authors would like to thank and acknowledge all collaborators from collected sites and animal owners for their participation in the study. In particular: Do Yew Tan, Na Lu, Yin Zhijuan, Jiangwei Wang, Xin Liu, Xinghui Chen, Dang Anh Thy, Junyan Dong, Isabelle Von Richthofen, Evonne Lim, Clair Chen, Mickael Banawa, Marielle Servonnet.

